# Front-end Weber-Fechner gain control enhances the fidelity of combinatorial odor coding

**DOI:** 10.1101/475103

**Authors:** Nirag Kadakia, Thierry Emonet

## Abstract

Odor identity is encoded by spatiotemporal patterns of activity in olfactory receptor neurons (ORNs). In natural environments, the intensity and timescales of odor signals can span several orders of magnitude, and odors can mix with one another, potentially scrambling the combinatorial code mapping neural activity to odor identity. Recent studies have shown that in *Drosophila melanogaster* the ORNs that express the olfactory co-receptor Orco scale their gain inversely with mean odor concentration according to the Weber-Fechner Law of psychophysics. Here we use a minimal biophysical model of signal transduction, ORN firing, and signal decoding to investigate the implications of this front-end scaling law for the neural representations of odor identity. We find that Weber-Fechner scaling enhances coding capacity and promotes the reconstruction of odor identity from dynamic odor signals, even in the presence of confounding background odors and rapid intensity fluctuations. We show that these enhancements are further aided by downstream transformations in the antennal lobe and mushroom body. Thus, despite the broad overlap between individual ORN tuning curves, a mechanism of front-end adaptation, when endowed with Weber-Fechner scaling, may play a vital role in preserving representations of odor identity in naturalistic odor landscapes.

---

Animals identify and discriminate odors using olfactory receptors (Ors) expressed in olfactory receptor neurons (ORNs) [1–4]. Individual ORNs, which typically express a single Or, respond to many odorants, while individual odorants activate many distinct ORNs [5–8]. Odors are thus encoded by the combinatorial patterns of activity they elicit in the sensing periphery [5–7, 9–11], patterns decoded downstream into behavioral response [12]. Still, ethologically-relevant odors are often mixed with background ones [13] and intensity can vary widely and rapidly as odors are carried by the wind [14–17]. How are odors recognized reliably despite these confounds? In *Drosophila melanogaster*, ORN dose response curves exhibit similar Hill coefficients but distinct power-law distributed activation thresholds [6, 18], which together with inhibitory odorants enhance coding capacity [6, 18–20]. In antennal lobe (AL) glomeruli, mutual lateral inhibition normalizes population response, reducing the dependency of activity patterns on odor concentration [21, 22]. Further downstream, sparse connectivity to the mushroom body (MB) helps maintain neural representations of odors, and facilitates compressed sensing and associative learning schemes [23–26]. Finally, temporal features of neural responses contribute to concentration-invariant representations of odor identity [27–30].

Here we examine how short-time ORN adaptation at the very front-end of the insect olfactory circuit contributes to the fidelity of odor encoding. Our theoretical study is motivated by the recent discovery of invariances in the response dynamics of ORNs expressing the co-receptor Orco. The responses of ORNs to diverse odorants of the same concentration differ widely, due to differences in odor-receptor affinities [6, 31, 32] and stimulus dynamics [33]. However, downstream from this input nonlinearity, signal transduction and adaptation dynamics exhibit a surprising degree of invariance with respect to odor-receptor identity: reverse-correlation analysis of ORN response to fluctuating stimuli produces highly stereotyped, concentration-invariant response filters [18, 33, 34].

These properties stem in part from an apparently invariant adaptive scaling law in ORNs: gain varies inversely with mean odor concentration according to the Weber-Fechner Law of psychophysics [35, 36], irrespective of the odor-receptor combination [34, 37, 38]. The invariance of this relatively fast adaptation (~250 ms) can be traced back to feedback mechanisms in odor transduction, upstream of ORN firing [34, 37–39], which depend on the activity of the signaling pathway rather than on the identity of its receptor [39]. The generality of the adaptive scaling suggests it could be mediated by the highly conserved Orco co-receptor [40–43]. Indeed, phosphorylation sites have been recently identified on Orco, some being implicated in odor desensitization, albeit over much longer timescales [43, 44].

While in a simple system such as *E. coli* chemotaxis [45], adaptive feedback via the Weber-Fechner Law robustly maintains sensitivity over concentration changes, the implication for a multiple-channel system – which combines information from hundreds of cells with overlapping receptive fields – is less straightforward. Here we combine a biophysical model of ORN adaptive response and neural firing with various sparse signal decoding frameworks to explore how ORN adaptation with Weber-Fechner scaling affects combinatorial coding and decoding of odor signals spanning varying degrees of intensity, molecular complexity, and temporal structure. We find that this front-end adaptive mechanism promotes the accurate discrimination of odor signals from backgrounds of varying molecular complexity, and aids other known mechanisms of neural processing in the olfactory circuit to maintain representations of odor identity across environmental changes.

## RESULTS

### Model of ORN sensing repertoire

We consider a repertoire of *M* = 50 Orco-expressing ORN types modeled using a simple extension of a minimal model of odor-to-ORN firing [34] that reproduces the type of Weber-Fechner adaptation and firing rate dynamics measured in individual *Drosophila* ORNs in response to Gaussian and naturalistic signals. Within ORNs of type *a* = 1*, …, M*, we model Or-Orco complexes as non-selective cation channels [40] (Fig. 1a) that stochastically switch between active and inactive states, while simultaneously binding to odorants *i* with dissociation constants, 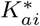 and *K*_*ai*_, respectively [34, 39]. Assuming these processes are faster than other reactions in the signaling pathway, the quasi-steady state active fraction *A_a_* of channels in ORNs of type *a* reads:

**Fig. 1:**
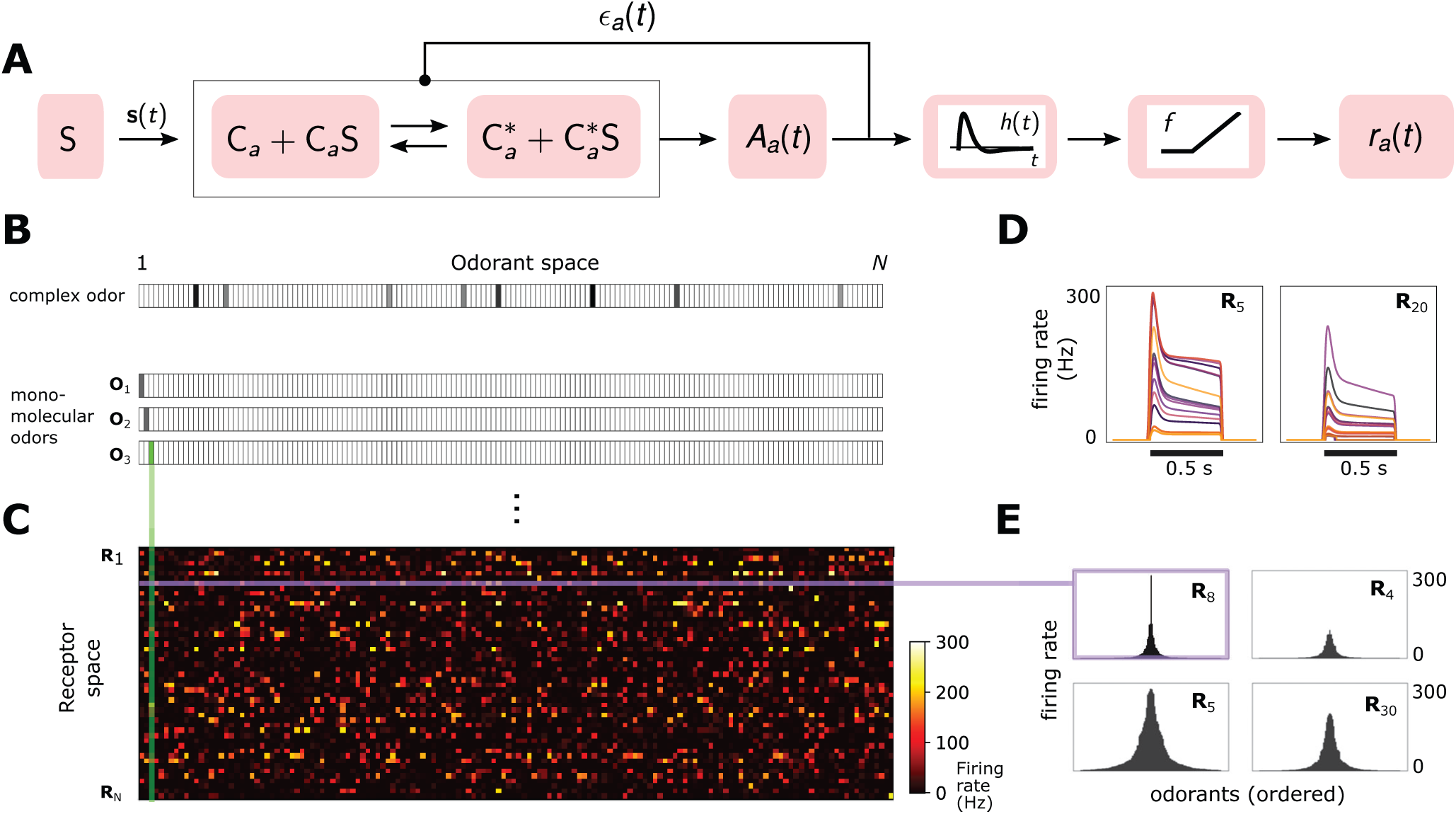
A Simple ORN model. Or/Orco complexes C_a_ respond to odor concentration *s(t) = (s_1_(t), …, s_i_(t), …, s_N_ (t))* by binding odorant molecules (S in the diagram) of type i and concentration s_i_(t), where t is time. Or/Orco complexes stochastically switch between active and inactive states, where the steady-state active fraction is determined by the free energy difference (in units of k_B_T) between active and inactive conformations in the unbound state, *ϵ_a_(t)*, and by odorant binding with dissociation constants 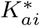 and *K* _ai_ (SI text). Adaptation is mediated by a negative feedback [39] from the activity of the channel onto the free energy difference *ϵ_a_(t)* with timescale τ. ORN firing rates *r_a_(t)* are generated by passing *A_a_(t)* through a linear temporal filter h(t) and a nonlinear thresholding function f. B Odor mixtures are represented by N-dimensional vectors s, whose components si are the concentrations of the individual molecular constituents of s. C Step-stimulus firing rate of 50 ORNs to the 150-possible monomolecular odors **s** = *s_i_*, given power-law distributed 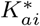 [18]. D Temporal responses of two representative ORNs to a pulse stimulus, for several monomolecular odorants (colors). E Representative ORN tuning curves (a single row of the response matrix in C, ordered by magnitude). Tuning curves are diverse, mimicking measured responses [6].

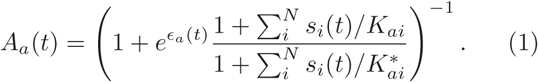

*s_i_*(*t*) are the time-dependent concentrations of the individual monomolecular components of the odor signal **s**(**t**) at time *t*, and *N* = 150 is the size of the molecular odorant space (Fig. 1b). Inward currents elicited by activating Or-Orco channels [40] eventually result in a negative feedback onto *A_a_*(*t*) [34, 38, 39]:

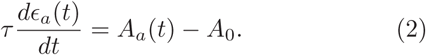

Here, *τ* is the adaptation time and *E_a_*(*t*) is the free energy difference in units of *k_B_T* between active and inactive conformations of the unbound Or-Orco channel. We assume changes in free energy are limited to the finite range *E*_L*,a*_ < E_*a*_(*t*) *< E*_H*,a*_ [34]. Firing rate is minimally modeled by filtering the activity *A_a_*(*t*) with the bi-lobed filter *h*(*t*) and rectifying nonlinearity *f* [34]:

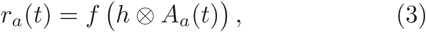

where⊗is convolution. When deconvolved from stimulus dynamics, the shapes of the temporal kernels of *Drosophila* ORNs that express Orco are largely receptor- and odor-independent [18, 33, 34]. Moreover, adaptation is not intrinsic to the receptor [39]. Accordingly, for simplicity *τ*, *A*_0_, *h*(*t*), and *f* are assumed independent of receptor and odorant identities.

We assume that the lower cutoffs *E*_L*,a*_ are receptordependent and choose them from a normal distribution. This variability ensures that ORNs are activated above quiescence (set at 5 Hz) at distinct stimulus levels [33, 34]. Diversity among odor-ORN responses arises mainly from the distribution of chemical dissociation constants (Fig. 1c). For simplicity we only consider agonists,i.e. 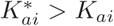, and assume receptors can only bind one odorant at a time. The analysis can easily be extended to include inhibitory odorants, which increases coding capacity [19]. We choose the dissociation constants from a power law distribution (*α* = 0.35) recently found across ORN-odor pairs in *Drosophila* larvae [18]. For a handful of ORNs we choose a very small value for one of the 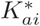 to mimic high responders to private odorants relevant to innate responses [32]. These private odors do not affect the general findings.

While this phenomenological model could be extended to include further details – e.g. we could relax the quasi-steady-state assumption in Eq. 1 and use a more complex model for neural firing [34] – this minimally-parameterized form captures the key dynamical properties of Orco-expressing ORNs relevant to our study: receptor-independent adaptation [39] with Weber-Fechner scaling [34, 37, 38] that maintains response time independent of mean stimulus intensity [33, 34], along with a diversity of both steady state and temporal firing patterns in response to a panel of monomolecular odorants [6, 27–29, 31] (Fig. 1d-1e).

### Concentration-invariant preservation of coding capacity and abstract representations of odor identity

To investigate how front-end Weber-Law adaptation might preserve representations of odor identity within the repertoire of ORN response, we project the 50dimensional firing rates **r** down to a 2-dimensional space using t-distributed stochastic neighbor embedding (tSNE) [46]. We first perform this embedding for an adaptive or non-adaptive system interacting with an odor environment containing a foreground odor A atop a background odor B (Fig. 2a). Both odors are sparse mixtures, with *K < N* odorants of similar concentrations, odor “identity” being the particular set of odorants in the mixture. Adaptation to the background is enacted by setting *E_a_* to their steady state values (via Eq. 2) in response to odor B alone. With adaptation in place, responses cluster by the identity of odor A, suggesting that ORN responses appropriately encode the identity of novel odors irrespective of background signals – once these backgrounds have been “adapted away” (Fig. 2a). This is notable, since, due to the combinatorics of ORN response (Fig. 1c), the adapted *E_a_* distributions are themselves heavily dependent on background identity. Responses in the non-adaptive system, meanwhile, exhibit no such clustering (Fig. 2a). A similar separation by odor identity is preserved in the adaptive system if we consider responses across different intensities (Fig. 2b).

**Fig. 2:**
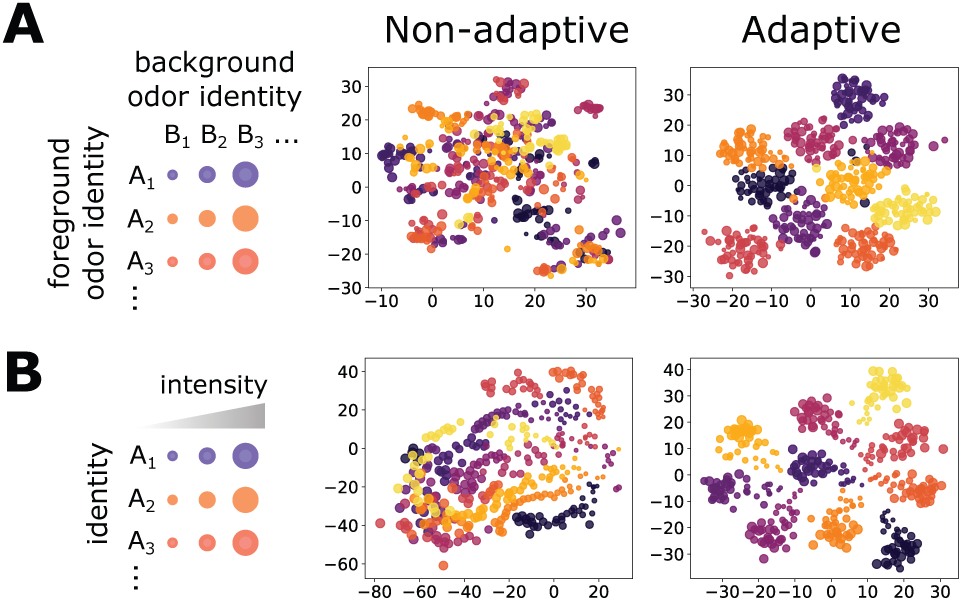
Front-end adaptation maintains representations of odor identity across background and intensity confounds. **A** Abstract representation of ORN responses in a low-dimensional embedding. Each point represents the repertoire of ORN firing rates, in response to an odor environment with both a foreground (point color) and background (point size) odor. In the adaptive system, Ea are set to their steady state values given odor B alone. **B** Similar to A, but now for odors whose concentrations span 4 decades (represented by point size). Here, the background odor identity is the same for all concentrations.

Preservation of these representations is enabled by the preservation of coding capacity. Accordingly, wecalculated the mutual information between odor and ORN response in time, verifying that the the adaptive system retains coding capacity as it confronts novel odors (Fig. S1). The non-adaptive system maintains coding capacity, though in a far more limited range of odor concentration.

### Front-end adaptation enhances odor discrimination in complex environments

How well does the preservation of coding capacity translate to better signal reconstruction? One potentially complicating factor is the disparity between sensor dimension and stimulus dimension: while *Drosophila* only express ~ 60 Or genes [47], the space of odorants is far greater [25]. However, many naturally-occurring odors are comprised of a small subset of odorants, which is suggestive as the theory of compressed sensing (CS) guarantees their reconstruction [48, 49]. It is unknown whether CS is implemented in the *Drosophila* olfactory circuit [50], and we use it mainly as a tool to quantify how front-end adaptation potentially affects odor decoding, later verifying our conclusions with other classification techniques that incorporate the known architecture of the olfactory system.

To incorporate the linear framework of CS, we treat the nonlinear odor encoding exactly but approximate the decoding to first order (this linearization simplifies the computation, but is not critical for our general results; see SI text and Fig. S7-S8). Odors **s** are assumed sparse, with *K ≪ N* nonzero components *s_i_* with mean concentration *s*_0_. We first examine how foreground odors are recognized when mixed with background odors of a distinct identity but similar intensities, quantifying decoding accuracy as the percentage of odors correctly decoded within some tolerance (Fig. 3a). Without adaptation, accuracy is maintained within the range of receptor sensitivity for monomolecular backgrounds, but is virtually eliminated as background complexity rises (Fig. 3b). The range of sensitivity is broader in the adaptive system, and is substantially more robust across odor concentration and complexity.

In realistic odor environments, the concentration and duration of individual odor whiffs vary widely [16]. We wondered how well a front-end adaptation mechanism with a single timescale *τ* could promote odor identity detection in such environments. As inputs to our coding/decoding framework, we apply a naturalistic stimulus intensity recorded using a photo-ionization detector [34] (Fig. 3c) to which we randomly assign sparse identities from the *N*-dimensional odorant space. To mimic background confounds, we combine these signals with static odor backgrounds, and then calculate the percentage of decoded whiffs. We assume the decoder has short-term memory: detected odor signals are only retained for *τ*_M_ seconds in the immediate past. Without ORN adaptation, sufficiently strong backgrounds eliminate the ability to reconstruct the identity of individual odor whiffs, irrespective of the complexity of either the foreground or background odor (Fig. 3d, blue lines). In the adaptive system, this is substantially mitigated (red lines in Fig. 3d), provided the memory duration *τ*_M_ is at least as long as the adaptation timescale *τ* (darker red lines). Because this short-term adaptation depends on the activity of the Or-Orco channel rather than on the identity of the receptor [33, 34, 39], the values of *τ* and *A*_0_ were assumed the same for all ORNs; still, our results hold if these invariances are relaxed (SI, Fig. S4-S5).

**Fig. 3:**
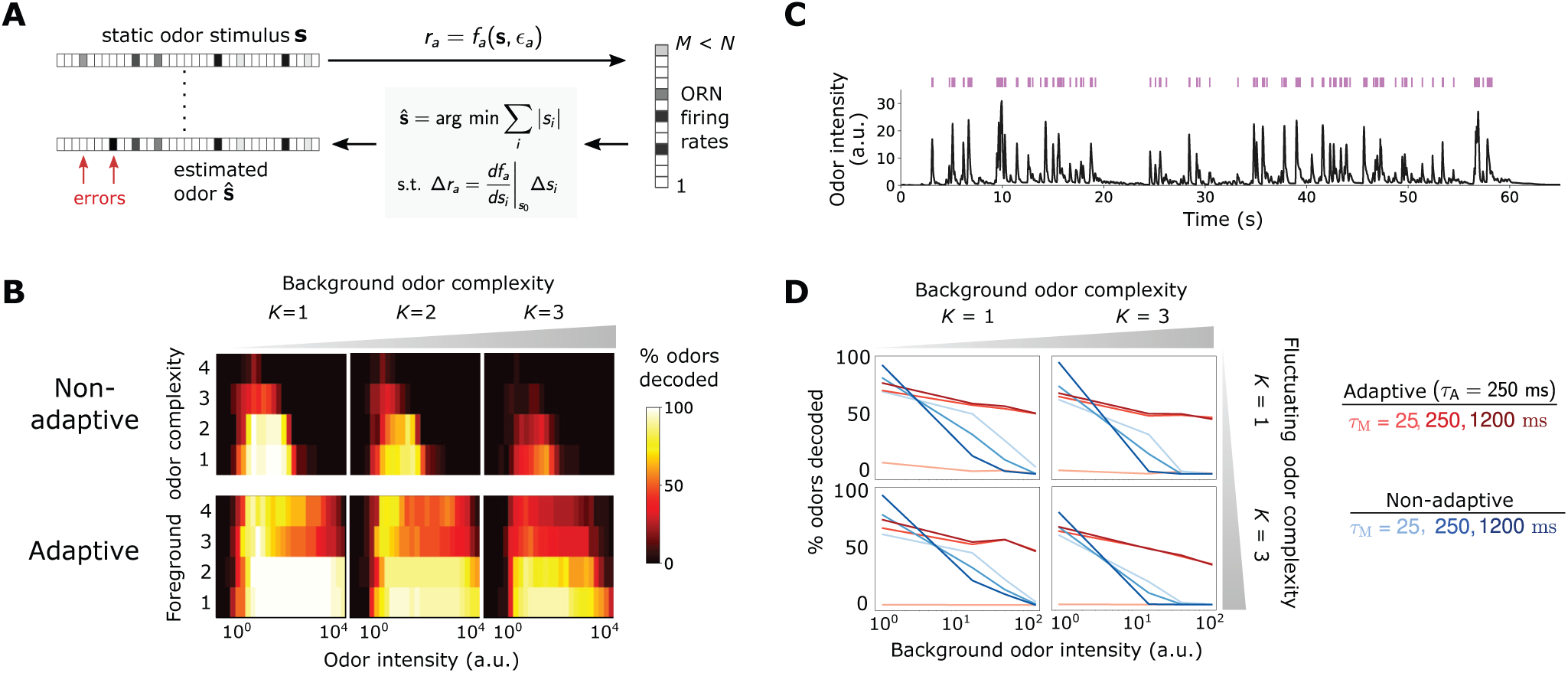
Front-end adaptation promotes accurate odor decoding in static and naturalistic odor environments. **A** Odor stimuli produce ORN responses via odor-binding and activation and firing machinery, as described by Eqs. 1-3. Odors are then decoded using compressed sensing by linearizing around a background s_0_ and minimizing the constrained *L_1_* norm of the odor signal. Odors are assumed sparse, with *K* nonzero components, *K ≪ N*. Odors are considered accurately decoded if the *K* sparse components are estimated within 25% and the *N-K* components not in the mixture are estimated below 10% of s_0_. *B* Decoding accuracy of foreground odors in the presence of background odors. *C* Recorded trace of naturalistic odor signal; whiffs (signal ¿ 4 a.u.) demarcated by purple bars. This signal is added to static backgrounds of different intensities and complexities. *D* Individual plots show the percent of accurately decoded odor whiffs as a function of background odor intensity, for the non-adaptive (blue) and adaptive (red) systems, for different t_M_ (line shades).

**Fig. 4:**
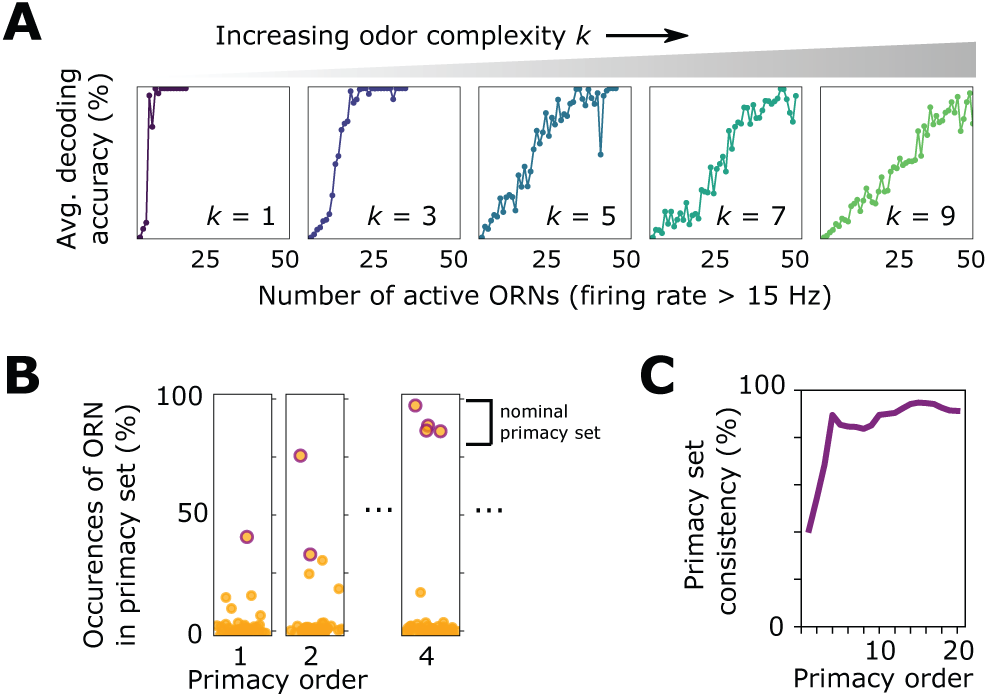
**A** Decoding accuracy as a function of the number of active ORNs, for different odor complexities. The primacy set consists of those ORNs required to be active for accurate decoding. **B** Frequency of particular ORNs in primacy sets of an odor placed atop different backgrounds. Individual plots show, for given primacy order *p*, the percentage of backgrounds for which the primacy set of odor A contains a given ORN (dots). Those with purple borders are the *p* most highly occurring – i.e. a nominal background-invariant primacy set for odor A. Points are jittered horizontally for visualization. **C** Consistency of primacy sets across backgrounds, as a function of *p*. Consistency is defined as the likelihood that an ORN in the nomimal primacy set appears in any of the individual background-dependent primacy sets, averaged over the nominal set (average of the y-values of the purple dots in B). 100% consistency means that for all backgrounds, the primacy set of odor A is always the same *p* ORNs.

### Front-end adaptation enhances primacy coding

The primacy coding hypothesis has recently emerged as an intriguing framework for combinatorial odor coding. Here, odor identity is encoded by the set (but not temporal order) of the *p* earliest responding glomeruli/ORN types, known as primacy set of order *p* [30]. If the activation order of ORNs were invariant to the strength of an odor step or pulse, primacy sets would in principle form concentration-invariant representation of odor identity. Though our coding framework uses the full ORN ensemble in signal reconstruction, some of these responses may contain redundant information, and a smaller primacy subset may suffice. To examine this, we apply our model to a sigmoidal stimulus that rises to half-max in 50 ms, calculating decoding accuracy in time. Since ORNs activate sequentially, the primacy set is defined by the ORN subset active when the odor is decoded. For simple odors, a limited set of earliest responding neurons fully accounts for the odor identity (Fig. 4a), in agreement with primacy coding. As expected for more complex odor mixtures, the full repertoire is required for accurate decoding. Primacy coding also predicts that for stronger stimuli, responses occur earlier, since the primacy set is realized quicker, which our framework replicates (Fig. S2).

Beyond mere consistency, however, front-end adaptation might also enhance primacy coding in different environments, such as background odors, which could scramble primacy sets. To investigate this, we considered again a sigmoidal odor step (odor A), now atop a static background (odor B) to which the system has adapted. We compared the primacy sets of odor A for 1000 different choices of odor B, finding that primacy sets are highly consistent across background confounds for all but the smallest primacy orders (Fig. 4b-4c). This also holds true for backgrounds of different concentrations (Fig. S2), suggesting a central role for front-end adaptation in reinforcing primacy codes across differing environmental conditions.

### Contribution of front-end adaptation for odor recognition within the *Drosophila* olfactory circuit

Signal transformations in the sensing periphery are propagated through the remainder of the olfactory circuit. How does front-end adaptation interact with these subsequent neural transformations? ORNs expressing the same OR converge to a unique AL glomerulus, where they receive lateral inhibition from other glomeruli [21, 51]. This inhibition implements a type of divisive gain control [22], normalizing the activity of output projections neurons, which then synapse onto a large number of Kenyon cells (KCs) in the mushroom body. To investigate how odor representations are affected by interactions between front-end ORN adaptation and this lateral inhibition and synaptic divergence, we extended our ORN encoding model by adding uniglomerular connections from ORNs to the antennal lobe, followed by sparse, divergent connections to 2500 KCs [23, 24, 52]. Inhibition was modeled via divisive normalization, with parameters chosen according to experiment [22]. We quantified decoding accuracy by training and testing a binary classifier on the KC activity output of sparse odors of distinct intensity and identity, randomly categorized as appetitive or aversive. For simplicity, odor signals of the same identity but differing intensity were assigned the same valence. We trained the classifier on *N*_ID_ sparse odor identities at intensities chosen randomly over 4 orders of magnitude, then tested the classifier accuracy on the same odor identities but of differing concentrations.

Classification accuracy degrades to chance level as *N*_ID_ becomes very large (Fig. 5a). When acting alone, either divisive normalization or ORN adaptation can help, although the effect of ORN adaptation is stronger. When both are active, accuracy improves further, suggesting that these distinct adaptive transformations may act jointly at different stages of neural processing in preserving representations of odor identity. As expected, these gains mostly vanish for the same odors chosen from a narrower range of concentrations (Fig. S3).

If we train the classifier to distinguish odors by identity rather than valence, the benefits conferred by divisive normalization do not appear until *N*_ID_ is substantial, with accuracy below 65% for *N*_ID_ *>* 50 (Fig. 5b). On the other hand, with ORN adaptation accuracy remains above 85% for more than 1000 odor identities, strongly implicating front-end adaptation as a key player in maintaining odor identity representations, before signals are further processed downstream.

**Fig. 5:**
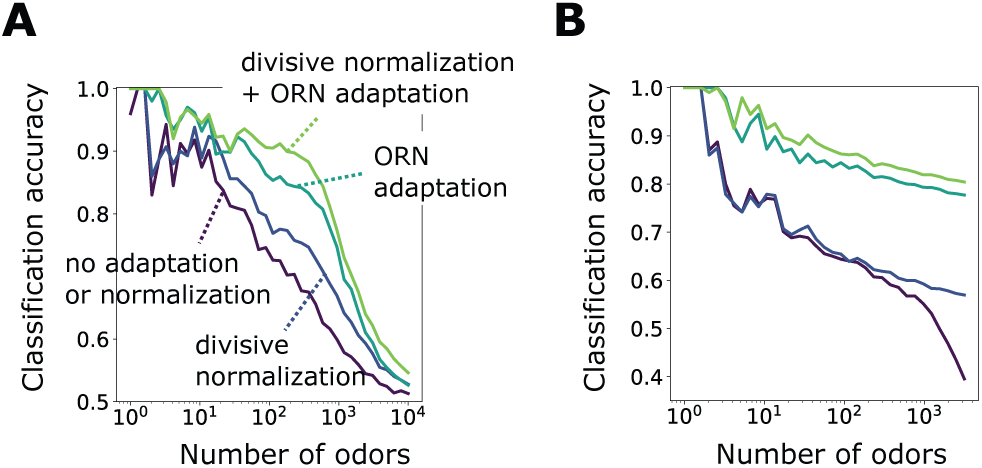
**A** Accuracy of binary classification by odor valence, as a function of the number of distinct odor identities classified by the trained network (concentrations span 4 orders of magnitude), in systems with only ORN adaptation, only divisive normalization, both or neither. **B** Same as (A) but now classifying odors by identity.

## DISCUSSION

We have found that Weber-Law adaptation at the very front-end of the insect olfactory circuit [34, 37, 38] may contribute significantly to the preservation of neural representations of odor identity amid confounding odors and intensity fluctuations. Drawing on experimental evidence for a number of ORN-invariant response features [18, 20, 33, 34, 39], we have found that this mechanism of dynamic adaptation confers significant benefits in coding fidelity, without the need for ORN-specific parameterizations. Still, our results hold when these invariances such as adaptation timescale or baseline activity are relaxed (SI Figs. S4-S5). In the olfactory periphery, front-end Weber Law adaptation therefore appears fairly robust, a consequence of controlling gain via feedback from channel activity [34, 39, 45], rather than through intrinsic, receptor-dependent mechanisms. While our framework incorporates many observed features of the *Drosphila* olfactory system – Weber-Law adaptation, power-law distributed receptor affinities, temporal filter invariance, connectivity topologies – it is minimal. We considered only one of the chemoreceptor families expressed in the fly antenna [1] and ignored possible contributions of odor binding proteins [53, 54], inhibitory odorants [19], and odorant-odorant antagonism [55], which could further boost coding capacity and preserve representation sparsity. Useful extensions to our nonlinear-linear-nonlinear model might incorporate ephaptic coupling between ORNs housed in the same sensillum [56], global inhibition in the mushroom body [57], and the effects of long-term adaptation [43].

Previous studies have characterized various neural mechanisms that help preserve combinatorial codes. Lateral inhibition between glomeruli helps tame saturation and boost weak signals [22]. The sparse degree of connectivity to either the olfactory bulb (vertebrates) or mushroom body (insects) may also be precisely tuned to optimize the capacity to learn associations [24]. In this work, we find that some of these downstream features act in concert with front-end dynamic adaptation in maintaining representations of odor identity.

Other studies have implicated the unique temporal patterns of neural response as signatures of odor identity [27–29, 58]. ORN and projection neuron time traces form distinct trajectories in low-dimensional projections, and cluster by odor identity, much as we have found here (Fig. 2). In our framework, temporal coding is implicit: because the input nonlinearity depends on the diversity of binding affinities, odor signals are naturally formatted into temporal patterns that are both odor-and ORN-specific (Figs. 1d-1e). Further, the short required memory timescales (*τ*_M_ ~ *τ* ~ 250 ms) suggest that only brief time windows are needed for accurate odor identification, consistent with previous findings [27]. Moreover, we find that front-end adaptation enhances the robustness of other combinatorial coding schemes, such as primacy coding [30], which relies on the temporal order of ORN activation but not absolute firing rate (Fig. 4).

In the well-characterized chemosensory system of bacterial chemotaxis, Weber Law adaptation is enacted through a feedback loop from the output activity of the receptor-kinase complexes onto the enzymes modifying receptor sensitivity [45]. It is interesting that some aspects of this logic are also present in ORNs: although the molecular players are different (and still largely unknown), it has been shown that transduction activity feeds back onto Or-Orco cation channel opening, enabling the Weber-Fechner relation [34, 38, 39]. That this adaptation mechanism appears to act similarly across ORNs [33, 34, 38] suggests the possible involvement of the universal co-receptor Orco, whose role in long-term adaptation has recently been reported [41–43]. Further, the identification of 4 subunits comprising the Orco-Or ion channel suggest that generic Or/Orco complexes may contain multiple odorant binding sites, which when included in our model supports our general findings (Fig. S6).

Weber Law ensures that sensory systems remain in the regime of maximum sensitivity, broadening dynamic range and maintaining information capacity [59]. For a single-channel system, this requires matching the midpoint of the dose-response curve to the mean ligand concentration [60], a strategy which may fail in multi-channel systems with overlapping tuning curves: adaptation to one signal could inhibit identification of others, if the signals excite some but not all of the same sensors. Our results show that this strategy is still largely functional. This can be traced to the observation that in CS, accuracy is guaranteed when sufficiently distinct odor identities produce sufficiently distinct ORN responses, a condition known as the restricted isometry property [49]. Indeed, the Weber-Fechner scaling increases the likelihood that this property is satisfied, beyond that in the non-adaptive system (SI text and Figs. S7-S8). Still, restricted isometry does not require that response repertoires are *invariant* to environmental changes. That is, even if the subset of active ORNs were concentrationdependent, odors could still in principle be fully reconstructible by CS. Nonetheless, our results in t-SNE clustering (Fig. 2), primacy coding (Fig. 4b-4c), and odor classification (Fig. 5) suggest that such response invariance is a natural byproduct of front-end adaptation. Together, this implies that Weber Law adaptation, whether required by the olfactory circuit for precise signal reconstruction (as in CS) or for developing odor associations (as in classification), can play an integral part in maintaining combinatorial codes amid changing environmental conditions.

## METHODS

Equations 1-2 are integrated numerically using the Euler method with a 2 ms time step. For ORN firing (Eq. 3), *h*(*t*) is bi-lobed [33]: *h*(*t*) = *Ap*_Gam_(*t*; *α*_1_*, τ*_1_) − *Bp*_Gam_(*t*; *α*_2_*, τ*_2_), *A* = 190, *B* = 1.33, *α*_1_ = 2, *α*_2_ = 3, *τ*_1_ = 0.012, and *τ*_2_ = 0.016, where *p*_Gam_ is the pdf of Gamma(*α*, 1*/τ*). Nonlinearity *f* is modeled as a linear rectifier with 5 Hz threshold. For t-SNE dimensionality reduction, ORN responses were generated for odor signal combinations consisting of 1 (among 10) distinct foreground odors A atop 1 (among 50) distinct background odors B, each of complexity *K* = 5, for Fig. 2a. Fig. 2b plots responses for 10 odors (*K* = 5) at 40 concentrations spanning 4 decades, atop a random sparse background odor (*K* = 5) of similar magnitude.

For compressed sensing decoding, sparse components *s_i_* are chosen as *s_i_* = *s*_0_+Δ*s_i_* where *s*_0_ is set as the center of linearization and Δ*s_i_ ∼ N* (*s*_0_*/*3*, s*_0_*/*9). Reconstructed signal components 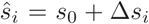 are computed by minimizing Σ*i |*Δ*s_i_|* subject to Δ*r_a_* = Σ*i dr_a_/ds_i_*|_*s0*_ Δ*s_i_* where Δ*r_a_* = *r_a_*(**S**) − *r*_*a*_(*s*_0_) are the “excess ORN firing rates about the linearization point. For static stimuli, *ϵ_a_*equals the fixed point of Eq. 2 in response to the background stimulus. For fluctuating stimuli, *ϵ_a_* is updated in time by continuously integrating *r_a_*(*t*), via Eqs. 2 and 3; thus, only knowledge of *r_a_*(*t*) is needed by the decoder. The naturalistic odor signal (Fig. 3d) was generated by randomly varying flow rates of ethyl acetate and measuring the concentration with a photo-ionization detector [34]. Statistics mirroring a turbulent flow [16] were verified (Fig. S9).

For the network model, the AL-to-MB connectivity matrix **J**_1_, is chosen such that each KC connects presynaptically to 7 randomly chosen AL glomeruli [23, 24]. The results shown in Fig. 5 are an average of 10 distinct instantiations of this random topology. The *Z* = 2500 KCs are then connected by a matrix **J**_2_ to a readout layer of dimension *Q*, where *Q* = 2 for binary and *Q* = *N*_ID_ for multi-class classification. Both AL-to-MB and MB-to-readout connections are perceptron-type with rectified linear thresholds. The weigh<u>t</u>s of **J**_1_ and **J**_2_ are chosen randomly from 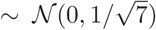 and 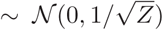, respectively. Only the **J**_2_ and the MB-to-output thresholds are updated during supervised network training, via logistic regression (for binary classification) or its higher dimensional generalization, the softmax cross entropy (for multi-class classification).

## ACKNOWLEDGEMENTS

NK was supported by a postdoctoral fellowship through the Swartz Foundation and TE by NIH R01 GM106189. We thank Damon Clark, John Carlson, Mahmut Demir, Srinivas Gorur-Shandilya, and Henry Mattingly for comments on the manuscript.

## Supporting Information Text

### Mathematical model

#### Model of odor binding, Or/Orco activation, and ORN firing

We model an odor as an *N*-dimensional vector **s** = [*s*_1_*, …, s_N_* ], where *s_i_ >* 0 are the concentrations of individual volatile molecules (odorants) comprising the odor. The olfactory sensory system is modeled as a collection of *M* distinct Or/Orco complexes indexed by the sub index *a* = 1, …, *M*, each of which can be bound with any one of the odorant molecules, and can be either active (firing) or inactive (quiescent). At first we assume there is one binding site per complex; this will be generalized to many sites. We consider the binding and activation processes to be in equilibrium, assigning each state a corresponding Boltzmann weight, where the zero of energy is set by the unbound, inactive state *C_a_*. These weights are:

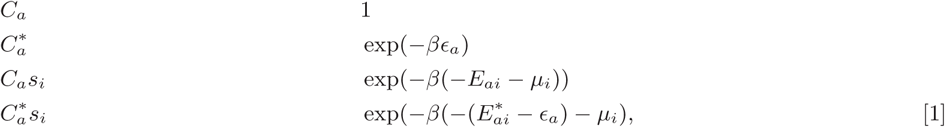

where *ϵ*_a_ (assumed positive) is the free energy difference between the active and inactive conformation of the unbound receptor, and *E_ai_* and 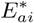 are the free energy differences (assumed positive) between the unbound and bound state for the inactive and active receptor, respectively. *μ_i_* = *μ*_0_ + *β*^−1^ log(*s_i_/s*_0_) is the chemical potential for odorant species *i* in terms of a reference chemical potential *µ*_0_ at concentration *s*_0_, *s*_0_ exp(−*βμ*_0_) = *s_i_* exp(−*βμ_i_*), which can be traded for the thermodynamic-relevant disassociation constants *K_ai_* = *s_0_e^β(−E_ai_−μ_0_)^*. Adding up contributions from all *i* odorants, the active fraction is:

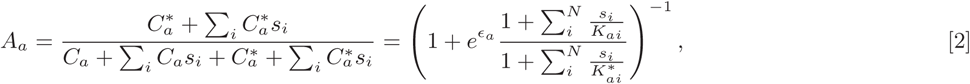

where we have expressed free energies in units of *k_B_T* = *β*^−1^ for notational convenience.

This expression can be generalized for the case of multiple, independent binding sites through some simple combinatorial factors. Consider first an odorant *i* which can bind one of two locations on receptor *a*. There are then 4 possible inactive states: both sites unbound, site 1 bound, site 2 bound, both sites bound. Combined with the active states, there are therefore 8 states for odorant *i* and receptor *a*, with energies:

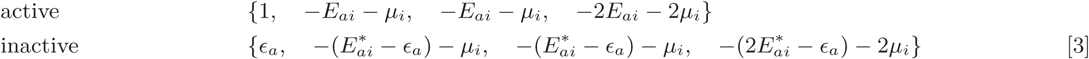

In the active fraction, Eq. 2, the Boltzmann factors combine through the binomial theorem, giving (for a single odorant environment *i*):

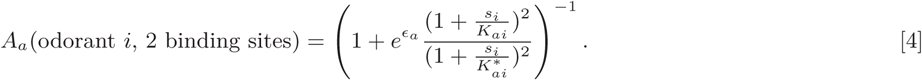

This expression generalizes for an arbitrary number of odorants and independent binding sites through the appropriate combinatorial factors, giving an active fraction of

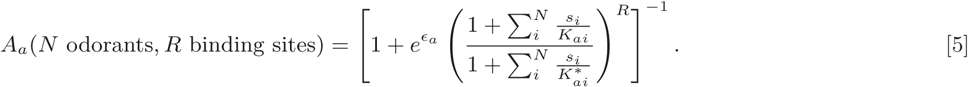

Finally, firing rate dynamics are assumed linear-nonlinear:

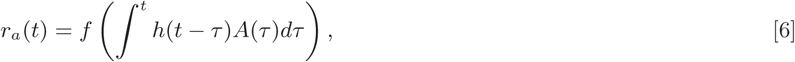

where *h*(*t*) and *f* are a temporal filter and rectifying linear unit (with threshold *θ* = 5 Hz) as noted in the main text.

### Compressed sensing decoding

#### Compressed sensing decoding of ORN response

Compressed sensing (CS) addresses the problem of determining a sparse signal from a set of linear measurements, when the number of measurements is less than the signal dimension. Specifically, it is a solution to

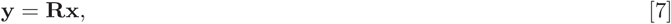

where **x** ∈ ℝ^*N*^ and **y** ∈ ℝ^*M*^ are vectors of signals and responses, respectively, and **R** is the measurement matrix. Since measurements are fewer than signal components, then *M < N*, whereby **R** is wide rectangular and so Eq. 7 cannot be simply inverted to produce **x**. The idea of CS is to utilize the knowledge that **x** is sparse, i.e.g only *K* of its components, *K ≪ N* are nonzero. Both the measurements and sparsity are thus combined into a single constrained optimization routine:

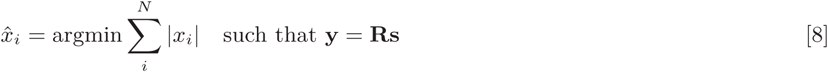

where 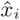 are the optimal estimates of the signal components and the sum, which is known as the *L*_1_ norm of **x**, is a natural metric of sparsity (1).

The *L*_1_ norm is a convex operation and the constraints are linear, so the optimization has a unique global minimum. To incorporate the nonlinear response of our encoding model into this linear framework, we assume that the responses are generated through the full nonlinear steady state response, Eq. 2- 6, but that the measurement matrix **R** needed for decoding uses a linear approximation of this transformation. Expanding Eq. 6 around **s**_0_ = **s** − Δ_**s**_ gives

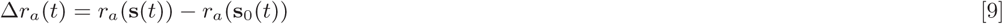

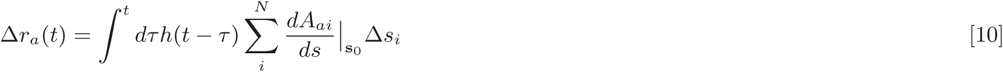

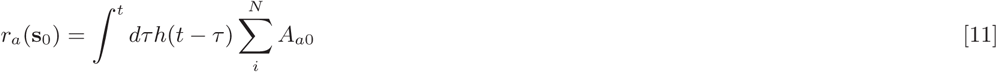

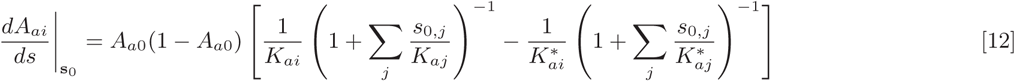

where *A*_*a0*_ = *A*(**s**_0_) and where Eqs. 10 and 11 hold only for integrands above 5 Hz (and are zero below), as per the linear rectifier *f*. We assume that the neural decoder has access to background **s**_0_, presumed learned (this assumption can be relaxed; see below), and to the linearized response matrix, Eq. 12, but must infer the excess signals Δ*s_i_* from excess ORN firing rates Δ*r_a_*(*t*). Thus, this corresponds to the CS framework (Eq. 8) via Δ**r** *→* **y**, Δ**s** *→* **x**, and 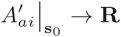. We optimize the cost function in Eq. 8 using sequential least squares programming, implemented in Python through using the scientific package SciPy.

#### Iterative Hard Tresholding (IHT) and the Restricted Isometry Property in compressed sensing

We stress that the purpose of response linearization is simply to apply compressed sensing reconstruction directly using linear programming, without worrying about issues of local minima in Eq. 8. This allows us to isolate the impact of Weber Law adaptation from the particularities of the numerics. An alternate technique for compressed signal reconstruction, *iterative hard thresholding* (IHT), does not minimize the constrained *L*_1_ norm directly, rather applying a hard threshold to an iteratively updated signal estimate (2). IHT can be generalized straightforwardly to nonlinear constraints, and would actually dispense with the need for a learned background **s**_0_, simply initializing the iterations from **s**_0_ = **0**. Remarkably, this technique works quite well even for non-linear measurements (3). We demonstrate the applicability of the IHT algorithm to our odor decoding system in Fig. S8, which reproduces qualitatively the findings in the main text. For these calculations, no background odor was assumed, each iterative decoding being initialized at the zero vector.

IHT provides an alternate computational technique of nonlinear CS, which could be used to both extend and verify our results. Further, it allows us to illustrate why Weber Law adaptation maintains signal reconstruction fidelity in our olfactory sensing model. Like CS using *L*_1_-norm minimization, IHT exhibits amenable reconstruction and convergence properties under the guarantee of the so-called restricted isometry property (RIP) (4). Loosely, RIP measures the closeness of matrix operator to an orthogonal transformation when acting on sparse vectors. The degree to which RIP is satisfied can be understood in terms of the spectrum of a measurement matrix **A**. In particular, if *λ_i_* are the eigenvalues of 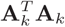, where **A**_*k*_ is any *k × m* submatrix of **A**, and

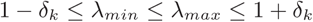

is satisfied for some *δ_k_*, then **A** satisfies the RIP with constant *δ_k_*. Plainly, the RIP states that the eigenvalues of 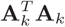, when acting on *k*-sparse vectors, are centered around 1. Thus, to intuit why signal reconstruction breaks down in the non-adaptive sensing system, we can investigate the eigendecomposition of various linearizations of the measurement matrix. We do this now, starting with a brief description of the IHT.

In the linear setting, IHT seeks sparse signals via the following iterative procedure (2):

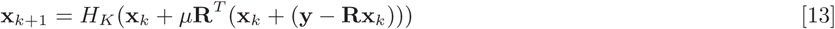

where **x**_*k*_ is the *k*th estimate of the sparse signal **x**, *μ* is a step size for the iterations, and **y**, **R** are as defined above. *H_k_*(⋅) is a thresholding function which sets all but the largest *K* values of its argument to zero. The nonlinear extension to IHT is (3):

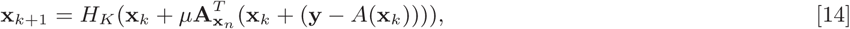

where *A* is a nonlinear sensing function and **A_x_**_*n*_ is a linearization of *A* about the point **x**_*n*_. Reconstructibility for *k*-sparse signals is guaranteed if **A_x_***n* satisfies RIP for all **x**_*n*_ and all *k*-sparse vectors (2). To get a sense of how this is preserved in the adaptive system, we calculate the eigenvalues for 1000 choices of **x**_n_, acting on random signals of given sparsity *K* (Fig. S7). Since the RIP is sensitive to constant scalings of the measurement matrix (while the actual estimation problem is not), we scaled all columns of **A_x_***_n_* to norm unity (5). This normalizes the eigenvalues of 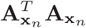 to center near unity before calculating the eigendecomposition, allowing us to assess the degree to which the RIP is satisfied. This scaled matrix can be used directly in Eq. 14 (3, 5). The spectra of these matrices indicates that the RIP becomes far more weakly satisifed in the non-adaptive system than in the adaptive one, for sufficient odor complexity and intensity.

---

**Fig. S1.**
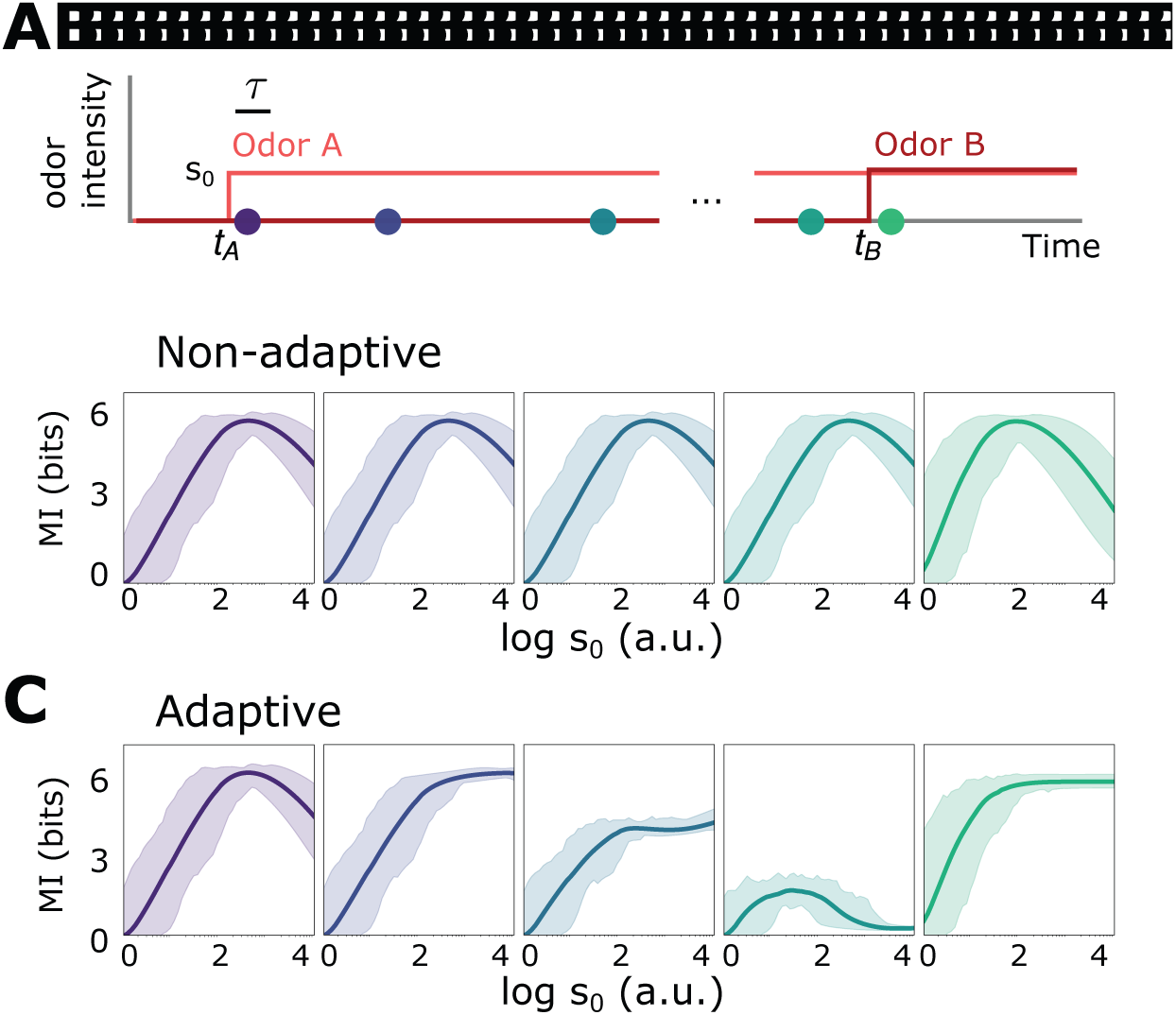
Front-end adaptive feedback preserves information capacity of the ORN sensing repertoire. Mutual information between signal **s**(*t*) = **s**_*A*_(*t*) + **s**_*B*_ (*t*) and response **r**(*t*) is calculated at various points in time *t* for an odor environment consisting of two step odors, A and B. **A** Odor A, with concentration **s**_*A*_(*t*), turns on at time *t_A_* and a odor B, with concentration **s**_*B*_ (*t*), turns on at some later time *t_B_*. Both odors have similar intensities *∼ s*_0_ and similar molecular complexity (*k* = 4). **B** Mutual information as a function of *s*_0_ for the non-adaptive system, respectively, at different time points after *t_A_*, corresponding to the dots in A. The mutual information carried by distinct ORNs is represented by the shaded region; their average is plotted by the heavy line. In the non-adaptive system, the mutual information peaks in the regime of high sensitivity after the arrival of odor A (purple, blue), and shifts leftward with the onset of odor B (teal, green). The leftward shifts occurs since stronger signals are more prone to response saturation (compromising information transfer) as odor B arrives. **C** Same as B, now for the adaptive system. The MI mimics the non-adaptive case at the onset of odor A, before adaptation has kicked in (purple). As the system adapts and responses decrease toward baseline, previously saturating signal intensities now cross the regime of maximal sensitivity, which therefore shifts rightward to higher *s*0 (dark blue). Much later, but before the arrival of odor B, the ORNs that responded now fire at a similar adapted firing rate *∼* 30 Hz, irrespective of odor identity, so the mutual information drops to zero. However, having now adjusted its sensitivity to the presence of odor A, the system can respond appropriately to odor B: the MI at *t_B_* is nearly 6 bits across decades of concentration immediately following *t_B_* (green).

**Fig. S2.**
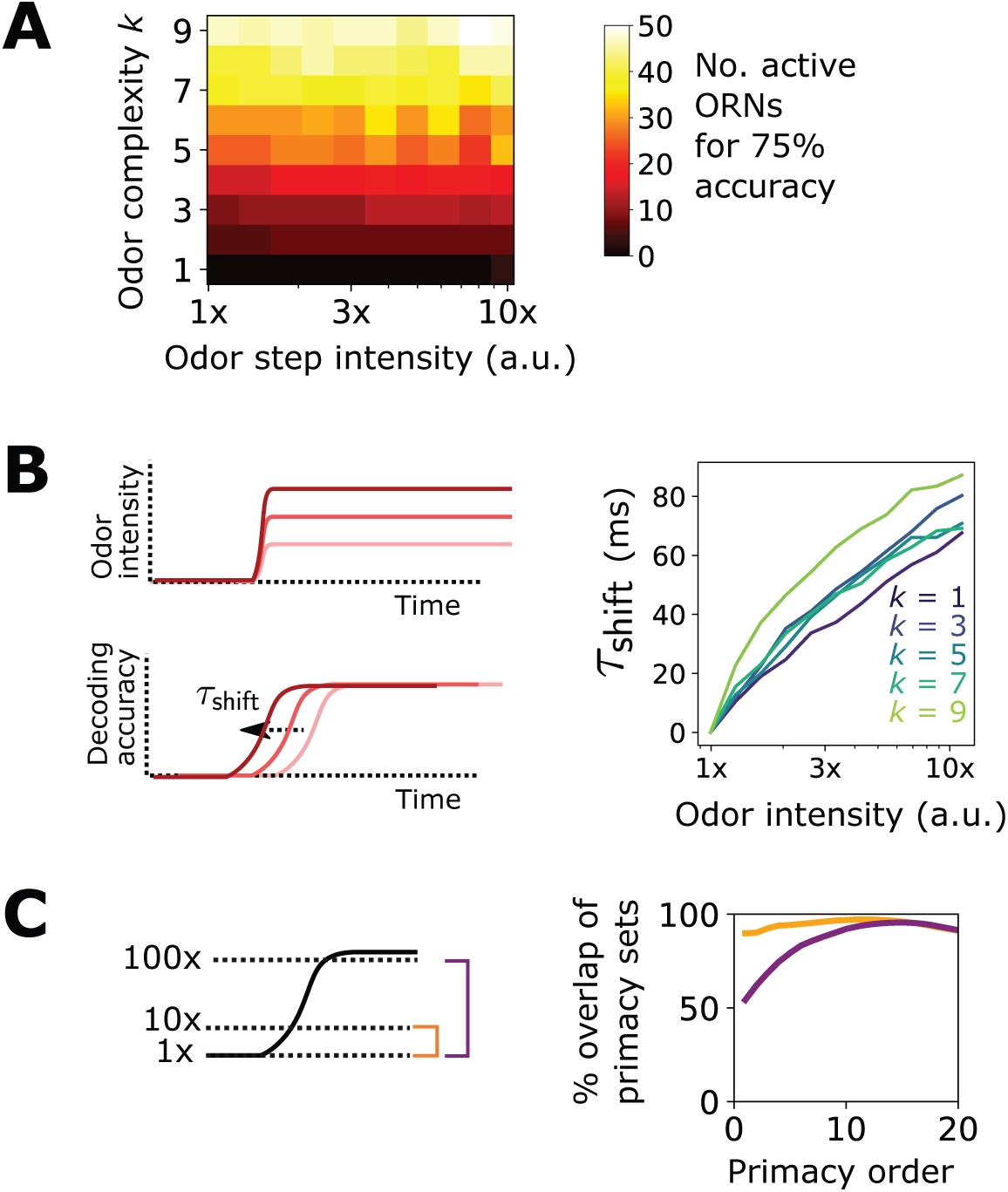
Additional results pertaining to the primacy coding hypothesis. **A** Percent of active ORNs required for 75% accuracy of a steep sigmoidal odor step, as a function of odor step intensity and odor complexity. For low complexities, a primacy set of fewer ORNs may be sufficient to decode the full odor signal; for higher complexities, the entire ORN repertoire is required. **B** In the primacy coding hypothesis, the primacy set is realized sooner for stronger odor signals, so odors are decoded earlier in time, resulting in a perceptual time shift with increasing odor concentration (6). We also find this shift in our compressed sensing decoding framework (right plot), which rises monotonically with step height for various odor complexities, in agreement with primacy coding. **C** The consistency of a primacy code across changes in background odor concentration, in a system with Weber Law adaptation. We calculate the primacy set for odor A (step odor; black) in the presence of either a weak, medium, or strong background (dotted lines; 1x, 10x, 100x a.u.), assuming the system has adapted its response to the background as described in the main text. Averaged across odor A identities, primacy sets for odor A when in the 1x background are nearly identical to those when odor A is in the 10x background (right plot; yellow). The same holds true when comparing the 1x and 100x backgrounds, for sufficiently large primacy order, above 8 or so right plot; purple). This indicates that Weber Law adaptation preserves primacy codes across disparate environmental conditions.

**Fig. S3.**
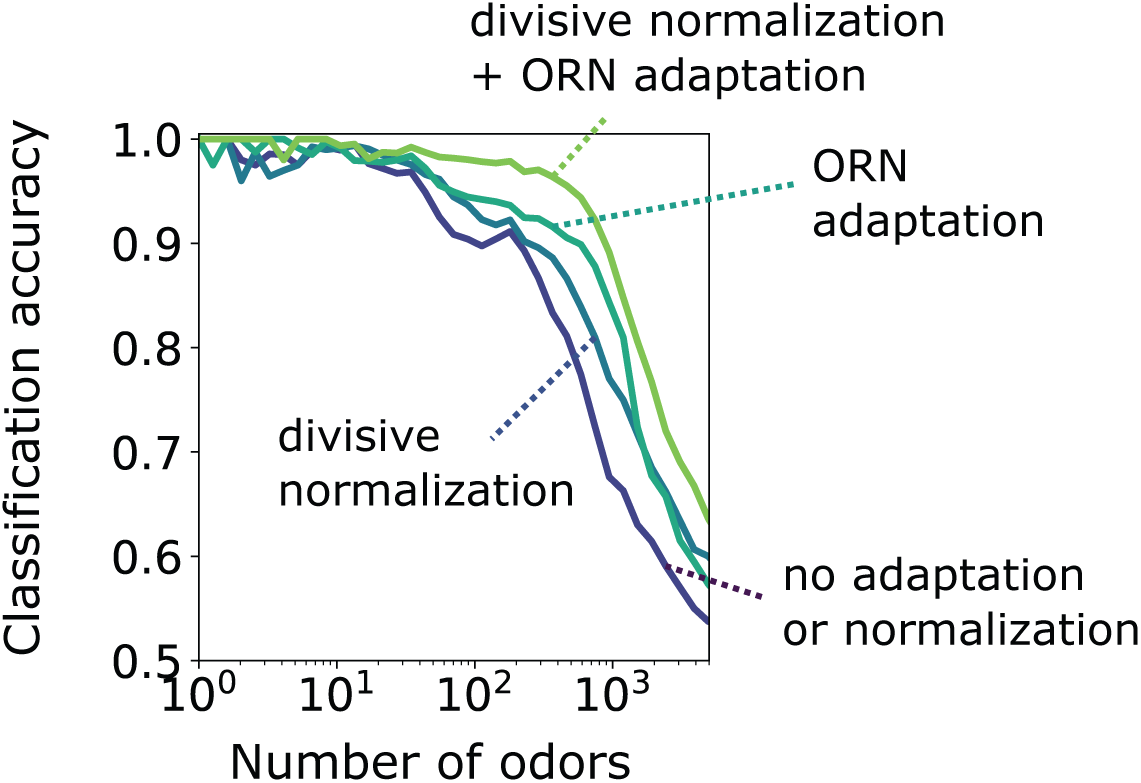
Accuracy of binary classification by odor valence, for odors whose concentrations span a narrow range of concentrations (1 order of magnitude). Accuracy is plotted as a function of the number of distinct odor identities classified by the trained network, in systems with only ORN adaptation, only divisive normalization, both or neither. Decoding gains conferred by divisive normalization and/or ORN adaptation are much smaller than when odors span a much larger range of concentrations, as shown in the main text.

**Fig. S4.**
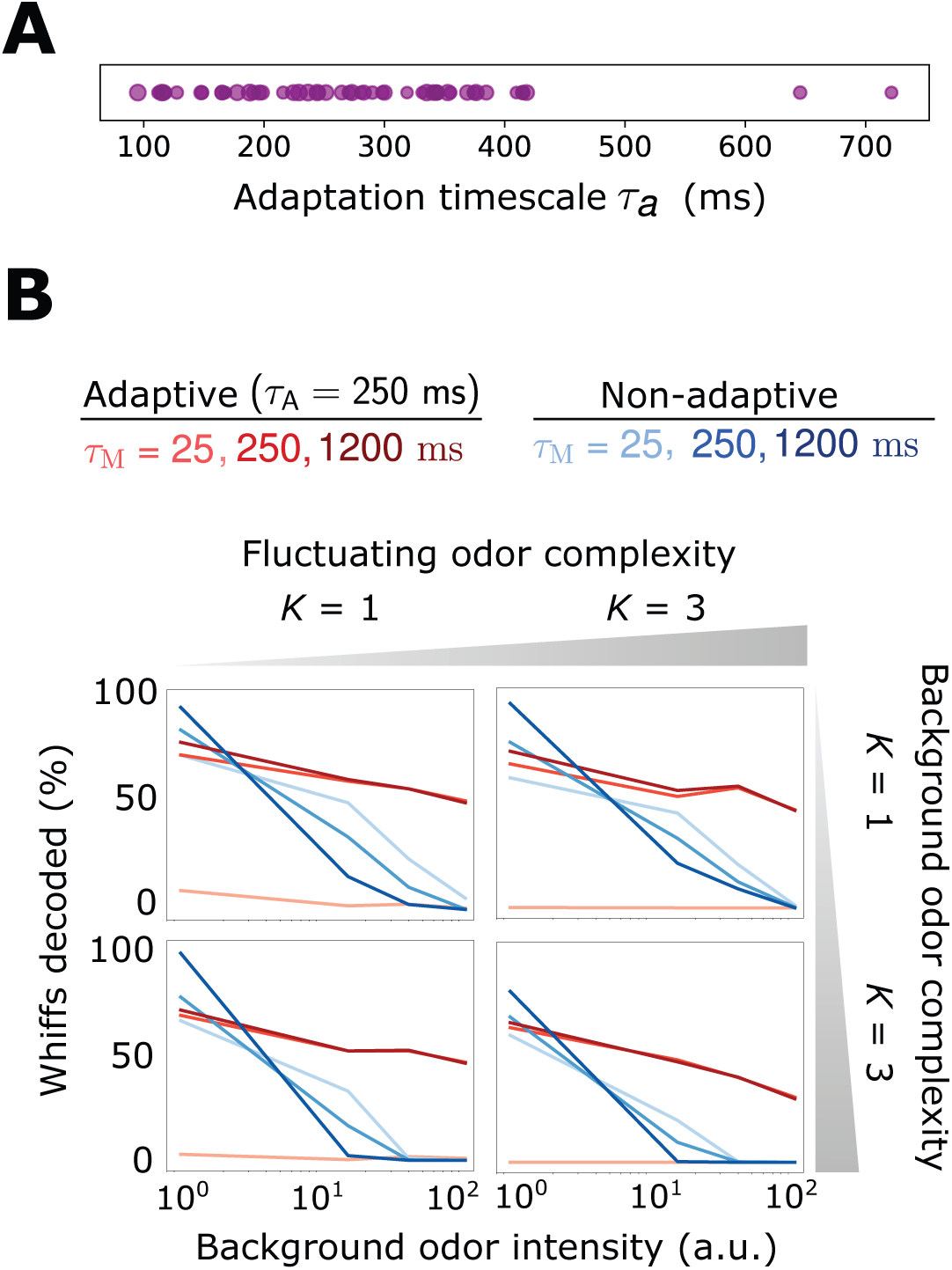
Decoding accuracy for system with broader distribution of adaptive timescales *τ*. **A** Distribution of timescales for all ORNs *a* (purple dots). Here, *τ_a_ ∼*= 10^*X*^ where *τ* = 250 ms as in the main text and *X ∼ N* (0, 0.2). **B** Individual plots show the percent of accurately decoded odor whiffs (same fluctuating odor signal used in the main text) as a function of background odor intensity, for the non-adaptive (blue) and adaptive (red) systems, for different *τ_M_* (line shades). Plots are arrayed by the complexity of the naturalistic signal (column-wise) and the complexity of the background odor (row-wise).

**Fig. S5.**
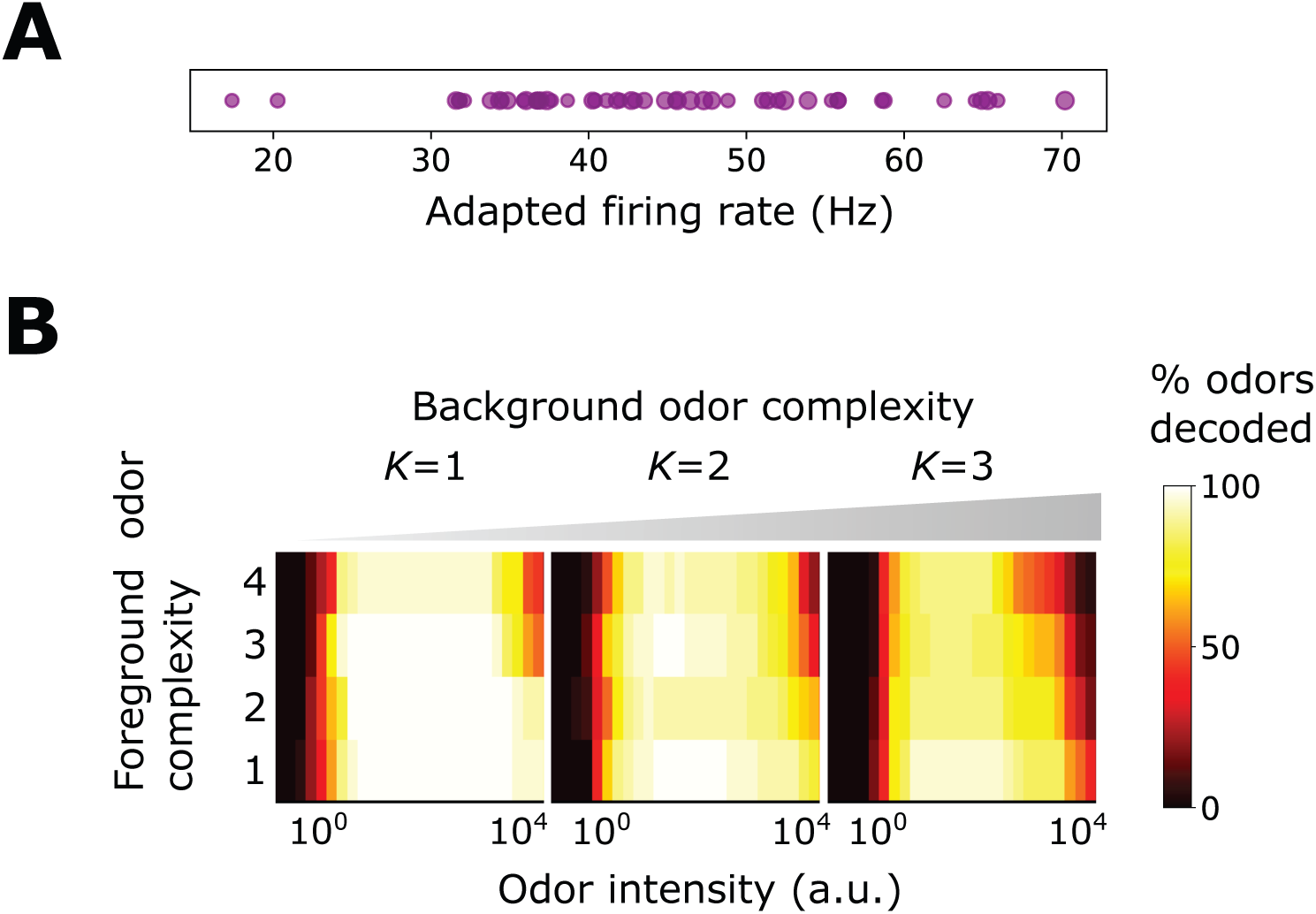
Benefits conferred by Weber-Fechner adaptation remain for a broader distribution of baseline firing rates *A*_a0_, now assumed to be ORN-dependent and chosen from a normal distribution. **A** Distribution of *A*_a0_. **B** Decoding accuracy of foreground odors in the presence of background odors.

**Fig. S6.**
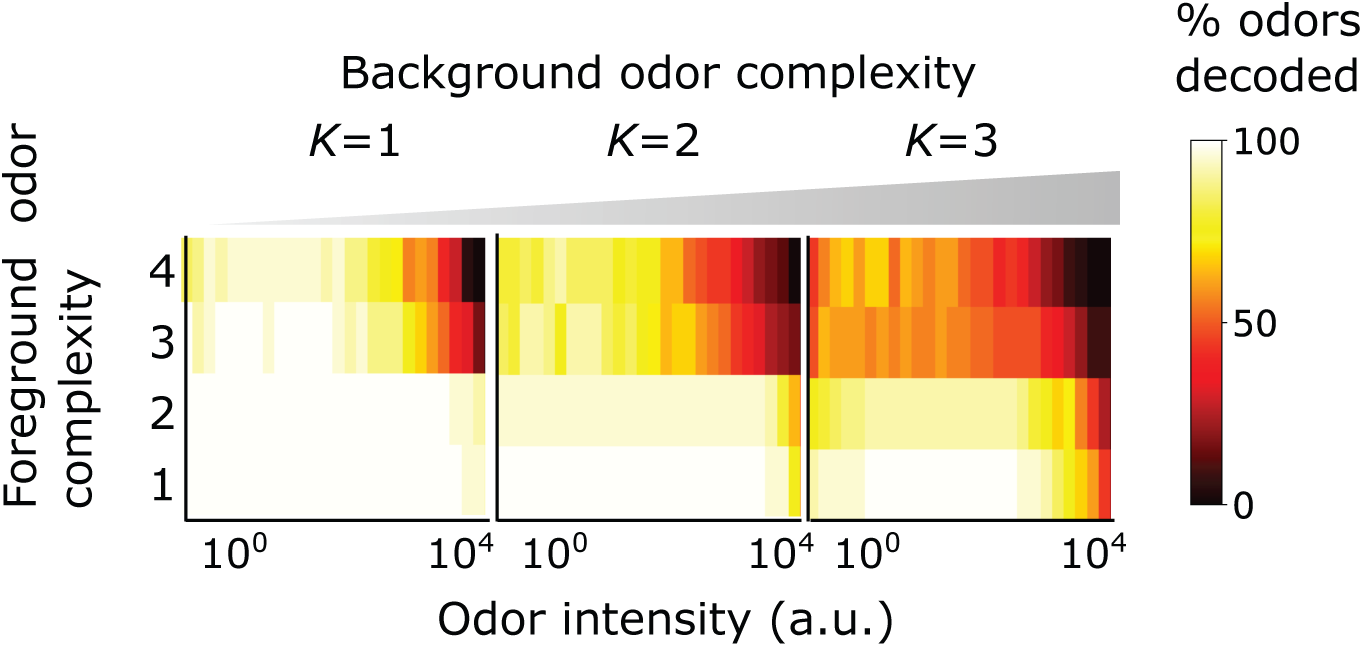
Benefits conferred by Weber-Fechner adaptation remain for 2 binding sites per receptor. This might conceivably occur in insect olfactory receptors, heterotetramers consisting of 4 Orco/Or subunits that gate a central ion channel pathway (7). Plotted is the decoding accuracy of foreground odors in the presence of background odors.

**Fig. S7.**
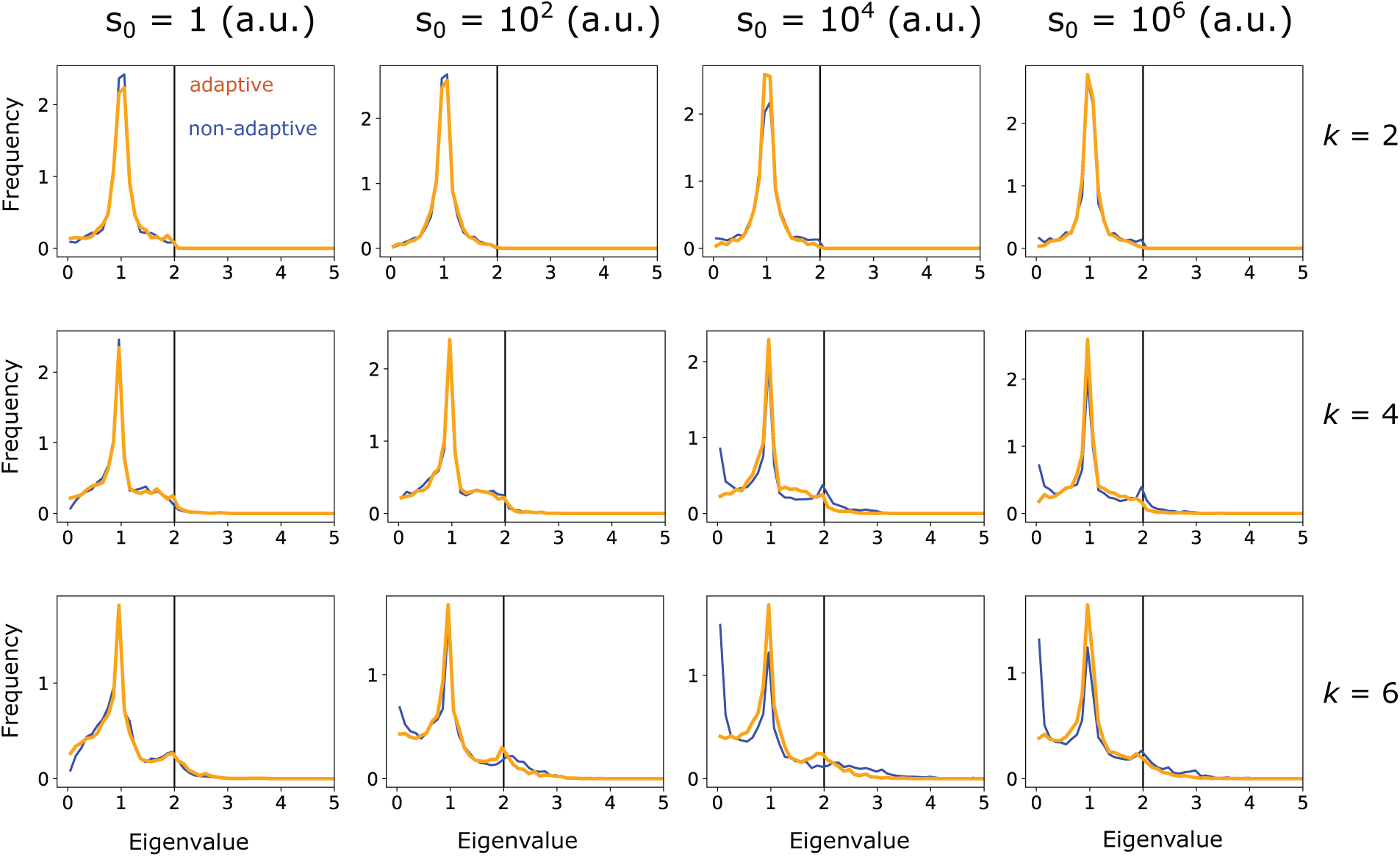
Eigenvalue distribution of 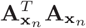, where **A_x_**_*n*_ is a *m × k* submatrix of the column-normalized linearized ORN response matrix **A**, evaluated at the linearization point **x***_n_*. Note that **x***_n_* is *k*-sparse, but its components do not necessarily align with the *k* columns chosen for the sub-matrix. Eigenvalues are calculated for the adaptive (orange) and non-adaptive (blue) systems, for 1000 randomly chosen linearization points **x***_n_* and submatrices. Plots are arranged for various odor sparsities (by row) and odor intensities (by column). The restricted isometry property is satisfied when the eigenvalues lie between 0 and 2 (black vertical line), and is more strongly satisfied the more centered the distribution is around unity. The increase in near-zero eigenvalues for the non-adaptive system at higher odor complexities and intensities (lower right plots) indicates the weaker fulfillment of the restricted isometry property for thhse signals, and leads to higher probability of failure in compressed sensing signal reconstruction.

**Fig. S8.**
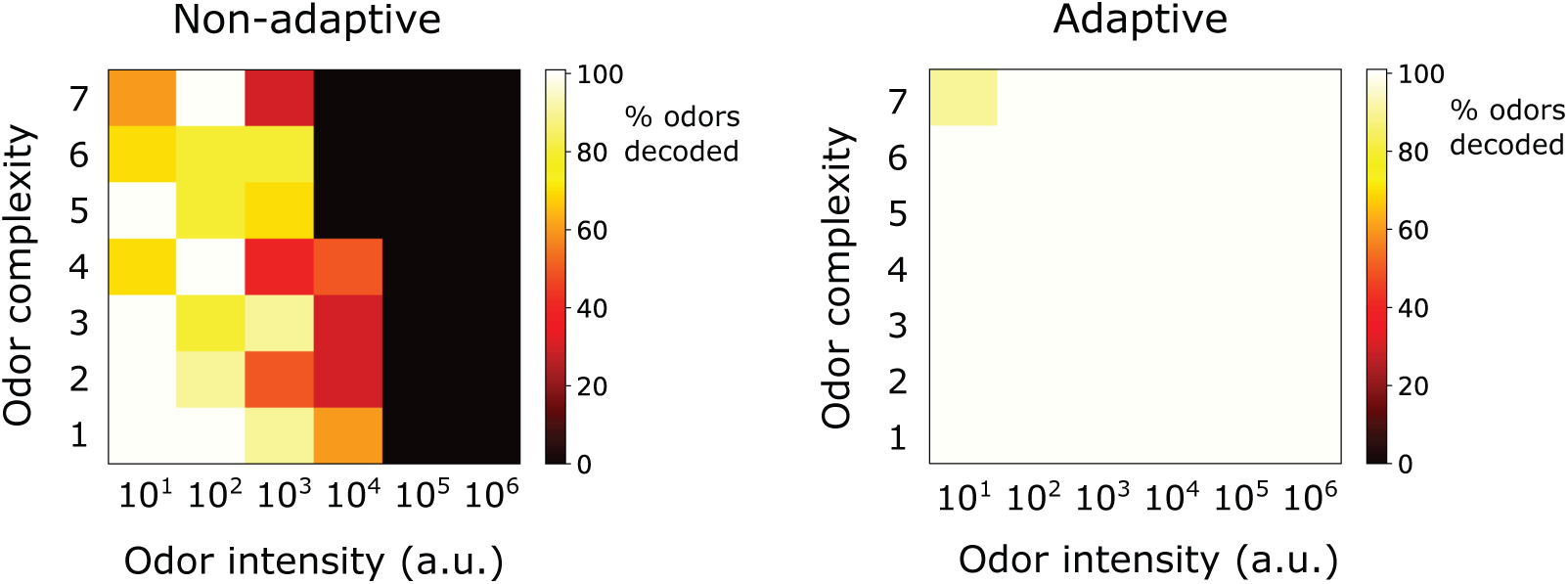
Decoding of odor signals (no background odors) using the IHT algorithm (2, 3) qualitatively reproduces the results from the main text, which used traditional CS with background linearization. In the adaptive case, IHT actually exhibits superior accuracy to traditional CS, though IHT demands more compute time. The results here show odor decoding accuracy for sparse odor signals of given complexity and intensity, averaged over 10 distinct identities. Odors are considered accurately decoded if the *K* sparse components are estimated within 25% and the components not in the mixture are estimated below 10% of *s*_0_. The iterative algorithm was initialized at 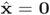 and run forward until 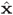 was stationary, or 10000 iterations were reached. Step size *μ* in Eq. 14 was set to *s*_0_*/*20. At each step, the linearized response (**A_x_**_*n*_ in Eq. 14) was evaluated at the result of the previous iteration. IHT also requires an assumption on the number of components in the mixture (which defines *H_K_* (⋅) in Eq. 14); here, that was set to twice the actual sparsity of true signal.

**Fig. S9.**
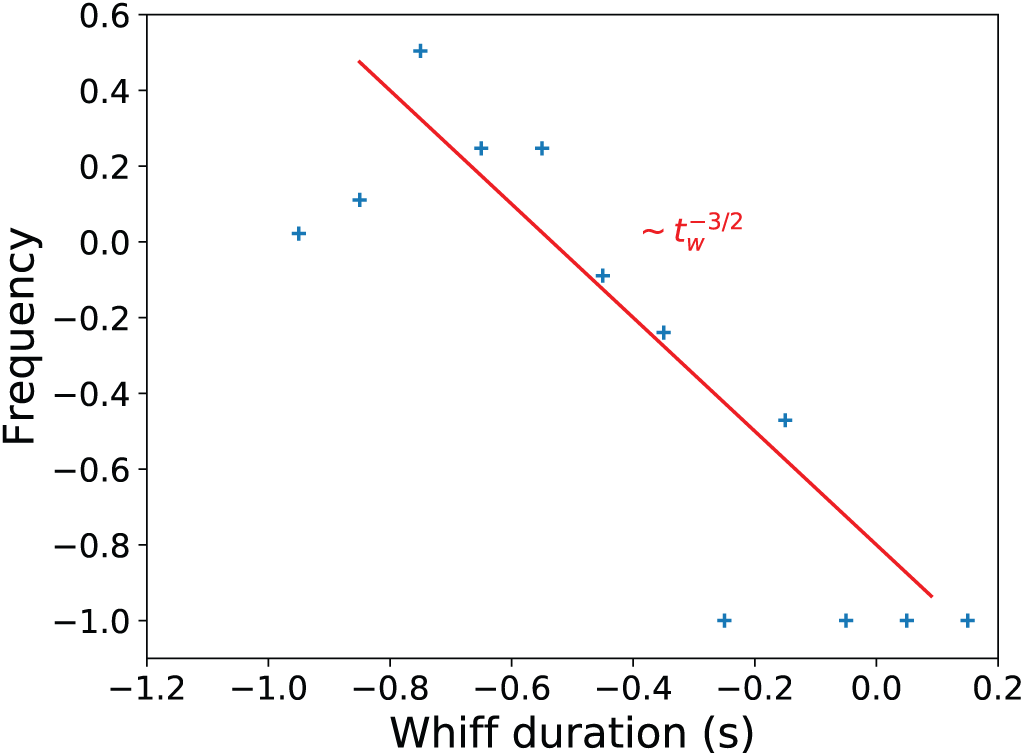
Distribution of whiff durations in naturalistic stimulus, compared to the theoretical prediction (8).

